# A globally diverse reference alignment and panel for imputation of mitochondrial DNA variants

**DOI:** 10.1101/649293

**Authors:** Tim W McInerney, Brian Fulton-Howard, Christopher Patterson, Devashi Paliwal, Lars S Jermiin, Hardip R Patel, Judy Pa, Russell H Swerdlow, Alison Goate, Simon Easteal, Shea J Andrews, for the Alzheimer’s Disease Neuroimaging Initiative

## Abstract

**Background:** Variation in mitochondrial DNA (mtDNA) identified by genotyping microarrays or by sequencing only hypervariable regions of the genome may be insufficient to reliably assign mitochondrial genomes to phylogenetic lineages or haplogroups. This lack of resolution can limit functional and clinical interpretation of a substantial body of existing mtDNA data. To address this limitation, we developed and evaluated a method for imputing missing mtDNA single nucleotide variants (mtSNVs) that uses a large reference alignment of complete mtDNA sequences. The method and reference alignment are combined into a pipeline, which we call MitoImpute.

**Results:** We aligned the sequences of 36,960 complete human mitochondrial genomes downloaded from GenBank, filtered and controlled for quality. These sequences were reformatted for use in imputation software, IMPUTE2. We assessed the imputation accuracy of MitoImpute by measuring haplogroup and genotype concordance in data from the 1,000 Genomes Project and the Alzheimer’s Disease Neuroimaging Initiative (ADNI). The mean improvement of haplogroup assignment in the 1,000 Genomes samples was 42.7% (Matthew’s correlation coefficient = 0.64). In the ADNI cohort, we imputed missing single nucleotide variants.

**Conclusions:** These results show that our reference alignment and panel can be used to impute missing mtSNVs in exiting data obtained from using microarrays, thereby broadening the scope of functional and clinical investigation of mtDNA. This improvement may be particularly useful in studies where participants have been recruited over time and mtDNA data obtained using different methods, enabling better integration of early data collected using less accurate methods with more recent sequence data.

## Background

Variation in mitochondrial DNA (mtDNA) is of interest because it is informative about human evolution [1] and because it is sometimes associated with disease [2]. Because human mitochondrial genomes do not recombine, the relationships among them can be described by a single phylogenetic tree. They can thus be grouped by the phylogenetic lineages to which they belong, into so-called haplogroups. In this system of evolutionary classification, genomes that belong to deeply divergent lineages form major haplogroups, with minor haplogroups corresponding to more recently diverged lineages.

In some studies, mtDNA is not fully characterised by whole genome sequencing, but rather by single nucleotide variants (mtSNVs) identified at predetermined sets of mitochondrial genome sites using microarrays. Partial mtSNV data obtained using such microarrays may be insufficient for reliable haplotype assignment of mitochondrial genomes. Reliable classification of mtSNV data is important because haplogroup classification is often used in population genetic studies and clinical investigations of associations between mitochondrial genomes and disease [3].

In addition, not all microarrays are designed to assay variation at the same sites in the human mitochondrial genome. Inconsistencies in the design of microarrays used in different studies can result in mtSNV datasets that are partially incompatible, making it difficult to combine them for joint analysis.

The duel problems of inaccurate haplotype assignment and incompatibility of data from studies that use different microarrays can be resolved by imputing mtSNVs at missing sites from a representative reference panel of human mitochondrial genome sequences. For incomplete mitochondrial genome sequence data, the base states (A, C, G, T) of missing nucleotide sites can be imputed by estimating their probabilities from the co-occurrence, as haplotypes, of bases at sites for which data are available. Accurate estimation of these probabilities has two fundamental requirements: (1) An accurate multiple sequence alignment (MSA) of genome sequences; and (2) A reference panel of genome sequences that is representative of the population being investigated.

The sequences of mitochondrial genomes vary substantially among human populations. To be representative, genome reference data must be obtained from the population that is the target of investigation. Data that is unrepresentative because it was obtained from an inappropriate population, can cause imputation to be biased and inaccurate [4–9]. Additional bias and inaccuracy may arise during construction of MSAs, which entails inserting alignment gaps (‘–’) between some of the nucleotides in some of the sequences being aligned – doing so accurately is a nontrivial challenge [10–12].

Imputation has been used to identify missing nuclear genome variants in incomplete sequence data using the 1,000 Genomes Project dataset [13, 14]. However, this dataset, which contains 2,504 nuclear and mitochondrial genome sequences representing 26 populations is only partially representative of human genome variation, with some populations (e.g., Pacific Islanders, Indigenous Australians, and Central Asians) still not represented.

In addition, considerable work is required to convert the 1000 Genomes Project mtSNV data from the format in which it has been made publicly available to a format that can be used for imputation. There is no published MSA of mitochondrial genomes from the 1000 Genome Project data or other more limited datasets (e.g., [3, 15]) that have been used for mitochondrial genome imputation. In addition to introducing errors, the need to recreate reference panels and MSAs for new studies results in a lack of the standardisation needed for comparison of results from different studies.

Imputation of mtDNA data would be greatly simplified and the substantial existing datasets of incomplete mitochondrial genome sequences would be made more accessible by overcoming the need for: preliminary data reformatting, identification and curation of suitable reference data panels, and standardisation of high-quality multi-sequence alignments.

Here we address these challenges by creating a large (n=36,960) globally diverse MSA using automated alignment software and manual curation by experienced researchers. This resource is publicly available as a standard reference panel on GitHub. We also describe a SnakeMake pipeline called MitoImpute, which we developed for easy imputation of mtSNVs through the IMPUTE2 framework [16]. Finally, we report our evaluation of MitoImpute using *in silico* microarrays (ISMs) derived from The 1000 Genomes Project Consortium [13] whole-genome sequence (WGS) data, and empirical data from the Alzheimer’s Disease Neuroimaging Initiative (ADNI) [17].

## Methods

### Reference Alignment and Reference Panel

Whole human mtDNA sequences were downloaded from GenBank on 18 July 2018 by adapting the MitoMap [18] search term (Supplementary Methods). This search returned 44,299 complete human mtDNA sequences and excluded archaic and ancient sequences (Table S1). Sequences were aligned to an pre-existing reference alignment (Supplementary Methods) in batches of 2,500 using MAFFT [19] using default settings in Geneious v10.2.6 [20]. The standardised site-numbering convention was maintained by including the revised Cambridge Reference Sequence (rCRS) [21] in both pre-existing and new reference MSAs, and by removing sites where gaps needed to be introduced to accommodate new sequences in the alignment.

To improve the quality of the MSA, sequences with ≥5 ambiguous characters or ≥8 gaps were removed. This threshold was set to enable the inclusion of haplogroup B sequences, which averaged 7 gaps relative to other sequences. This quality filter reduced the Reference Panel to 36,960 sequences (Table S1). To avoid adding bias to population frequency estimates, GenBank accessions with identical sequences were retained on the basis that they represent relatively common mitochondrial genomes.

AliStat v1.11 [22] was used to quantify the completeness of the new Reference MSA. The Reference Panel was created by converting the Reference MSA to formats compatible with IMPUTE2 [16].

### Validation Panel

In silico microarrays (‘microarray’ datasets) were created by selecting only mtSNVs present in commercially available microarrays from the 1000 Genomes Project Phase 3 WGS data (n=2,535). Microarray information was obtained from strand orientation files available from the Wellcome Centre for Human Genetics at the University of Oxford [23], with 103 strand files containing mtSNVs (Table S2). Haplogroup assignment for the WGS data and the ISMs was performed using HaploGrep2 [24] and Hi-MC [25].

### Imputation

We used the IMPUTE2 chromosome X imputation protocol [3, 16] to generated ‘imputed’ datasets from the microarray datasets and the reference panel. No recombination was assumed (i.e., a uniform recombination rate of r=0 across all sites). The Markov chain Monte Carlo step in IMPUTE2, which is used to account for phase uncertainty in recombining diploid data [16], was not used because human mitochondrial genomes are haploid and are not known to recombine. Only high-quality imputed sites were retained by removing sites with an IMPUTE2 information score of ≤ 0.3.

The effect of varying the number of sequences in the reference alignment (k_hap_) was estimated by setting k_hap_ to 100, 250, 500, 1,000, 2,500, 5,000, 10,000, 20,000, and 30,000. We tested the ability of our pipeline to impute rare variants by filtering the Reference Panel to exclude variants with minor allele frequencies (MAF) of MAF>1%, MAF>0.5% and MAF>0.1%, resulting in 409, 682 and 1874 mtSNVs, respectively (Table S3). With this filtering scheme, 2 of the 103 strand files did not include any mtSNVs at MAF>1% or MAF>0.5% (Table S2). Imputation accuracy was assessed using Matthews [26] Correlation Coefficient (MCC) for genotype concordance. Both HaploGrep2 [24] and Hi-MC [25] were used for haplogroup assignment, with the WGS data used as the truth set. HaploGrep2 has the advantage of covering the full scope of the PhyloTree haplogroup nomenclature [24, 27], including small sub-haplogroups. Hi-MC was developed for epidemiological research that uses high-throughput data by reducing PhyloTree nomenclature to 46 common haplogroups using a limited array of mtSNVs from which to assign haplogroups. We treated the first major sub-haplogroup of all L linages (i.e. L0), as well as HV and JT as macrohaplogroups.

Linear mixed-model ANOVA was used to assess the meaningfulness of difference in MCC (mean of mtSNVs per ISM) and haplogroup assignment for different parameters tested for k_hap_ and MAF.

Pipelines for implementing our imputation protocol and reproducing our results were initially created in BASH shell scripts then lifted over into SnakeMake [28] for the MitoImpute pipeline.

## Results

### Reference Alignment and Reference Panel

To comply with minimum reporting standards for MSAs, completeness metrics of the Reference MSA were computed (Table 1). As described in Wong, Kalyaanamoorthy [22], *C*_*a*_ is the completeness of the MSA, *C*_*r*_ is the completeness of the *r*^th^ sequence, *C*_*c*_ is the completeness of the c^th^ site, and *C*_*ij*_ is the completeness of the *i*^th^ and *j*^th^ sequences. Overall, the Reference MSA is highly complete (*C*_a_ > 0.99). Individual sequences are also mostly complete (*C*_*r*_), with the least complete sequence containing completely-specified nucleotides at 91% of its sites and the most complete sequence containing completely-specified nucleotides at all of its sites. The least complete site in the MSA contained completely-specified nucleotides in 44.3% of sequences, and the most complete sites had completely-specified nucleotides in all of the sequences. The proportion of homologous sites with completely-specified nucleotides at in both sequences (*C*_*ij*_) ranged from 83% so 100%, suggesting that the majority of sequence pairs contain enough information to quantify evolutionary distances. Sites and sequences missing a substantial number of nucleotide states were removed in the filtration processes as described in the Methods section.

**Table 1.** AliStat completeness metrics for the Reference MSA.

| Feature | Value(s) |
| --- | --- |
| Sequences | 44,299 |
| Sites | 16,569 |
| Completeness Score ($C_a$) | 0.9997 |
| C-score for individual sequences ($C_r$) [min-max] | 0.9119 - 1.0000 |
| C-score for individual sites ($C_c$) [min-max] | 0.4429 - 1.0000 |
| C-score for pairs of sequences ($C_{ij}, i \neq j$) [min-max] | 0.8314 - 1.0000 |

$C_a$ : completeness of the alignment; $C_r$ : completeness of the $r^{\text{th}}$ sequence; $C_c$ : completeness of the $c^{\text{th}}$ site; and $C_{ij}$ : completeness of the $i^{\text{th}}$ and $j^{\text{th}}$ sequences.

GenBank metadata on geographic provenance was available for 7,128 (19.3% filtered and 16.1% unfiltered) sequences in the Reference Panel, from 49 countries and 54 sub-country regions (Table S4). These regions included smaller ethnic groups such as Yami Taiwanese, Moroccan Berbers, Pacific Islanders, Indigenous Australians, and people from Central Asia and Siberia. For sequences with provenance information, there is, however, a distinct bias towards Europe (3,855; 54.1%; 10.4% filtered; 8.7% unfiltered) and East Asia (2,065; 29.0%; 5.6% filtered; 4.7% unfiltered).

All major haplogroups are represented in the Reference Panel (Figure 1, Table S1), including rare haplogroups such as haplogroup S, which is endemic to Indigenous Australians, haplogroup L5, which is found in Mbuti Pygmies, haplogroup L6, which is found in low frequencies in Yemen and Ethiopia, and haplogroups O and Q, which are found exclusively in the Pacific Islands. Haplogroup B was the haplogroup most frequently removed by the quality control filter (3,395 or 46% of all 7,339 removed sequences), leaving only 273 haplogroup B sequences. Haplogroup H was also heavily filtered following quality control (1,376; 19%), but remained well represented in the final reference panel (n=7,644). Only a small fraction of other haplogroups were removed during quality control.

**Figure 1:**
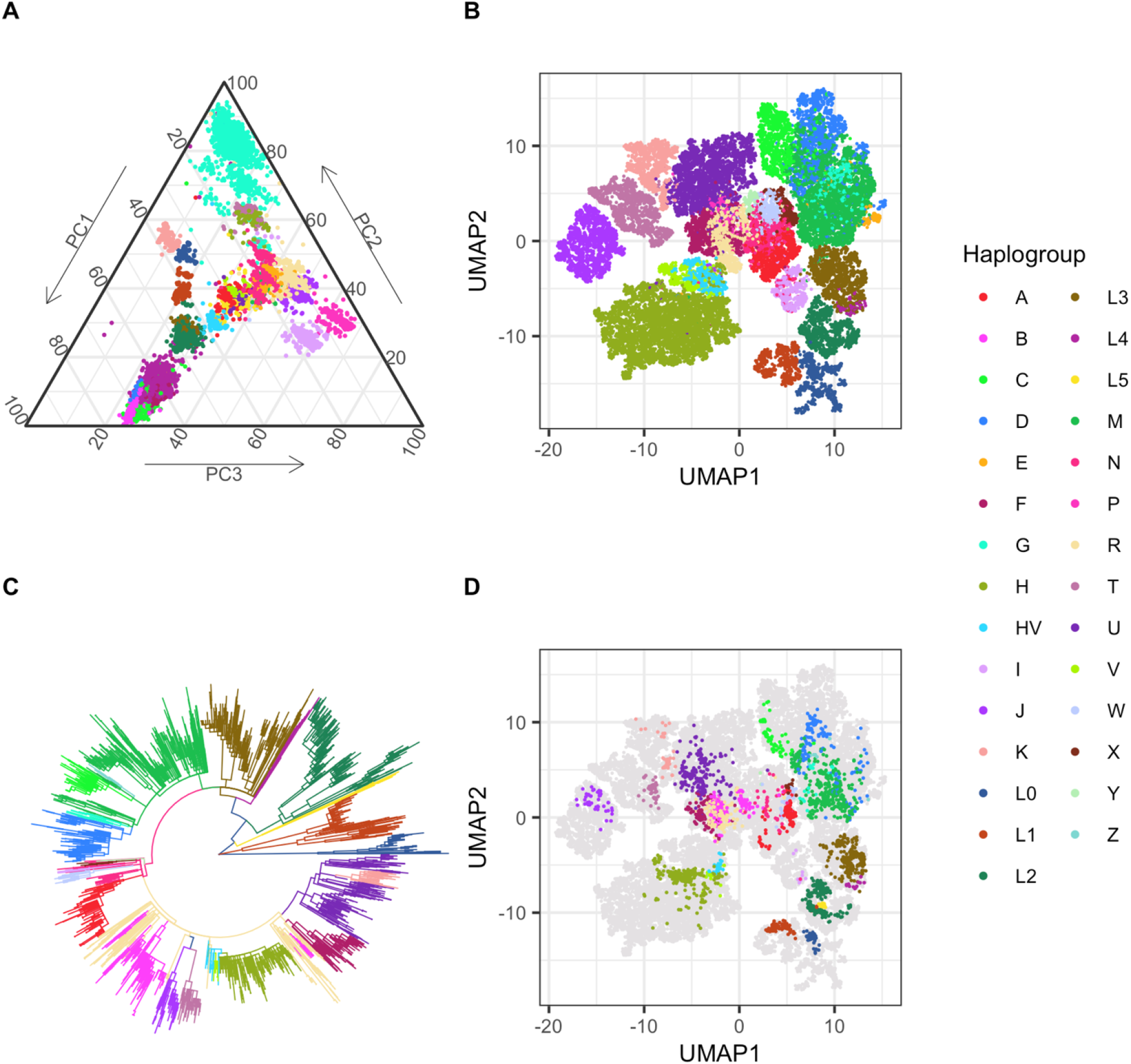
Diversity of mitochondrial reference alignment. A) PCA of mitochondrial sequencies included in reference panel coloured by haplogroup. B) UMAP projection of mitochondrial sequencies. C) Phylogenetic tree of 1000 genomes mitochondrial sequences highlighting phylogenetic relationship between mitochondrial haplogroups. D) Projection of 1000 Genomes mitochondrial sequences onto the mitochondrial reference alignment.

### *In silico* Microarrays

#### Parameter Tuning

We measured imputation accuracy of genotypes using the Matthews [26] Correlation Coefficient (MCC). To summarise MCC values, we calculated the mean MCC across all imputed sites, then compared the estimated marginal means using a linear mixed-model ANOVA. Our results show that the Reference Panel filtered to MAF>0.1% was the best performing (μ_*MCC*_ = 0.60), followed by MAF>0.5 (μ_*MCC*_ = 0.58), then by MAF>1% (μ_*MCC*_ = 0.57). These contrasts are all statistically significant (ANOVA, *p* = 0.002) (Table S5a-c). For the *k*_*hap*_ parameter, there was no significant pairwise differences between *k*_*hap*_ = 100 and the other *k*_*hap*_ values up to 1,000. Above a *k*_*hap*_ = 1,000, contrasts were often statistically significant (Table S5d-f), with larger *k*_*hap*_ parameter values performing comparatively poorly, indicating a reduced ability to correctly assign haplogroups for some ISMs.

Imputation accuracy was also evaluated using the IMPUTE2 Info Score using the same statistical framework described for MCC. In contrast to MCC, the Reference Panel filtered to MAF>1% was the best performing (μ_info_ = 0.73), followed by MAF>0.5 (μ_*MCC*_ = 0.69), and MAF>0.1% (μ_*MCC*_ = 0.63). All of these contrasts are statistically significant (ANOVA, *p* < 0.0001) (Table S6a-c). Starting at *k*_*hap*_ = 1,000, pairwise comparison of larger k_hap_ values become statistically significant, suggesting a meaningful difference in mean haplogroup concordance becomes apparent when more reference haplotypes are included.

Imputation accuracy was further evaluated by determining whether haplogroup assignments were concordant between imputed sequenced datasets. As HaploGrep2 assigns haplotypes to very specific sub-haplogroups, we measured concordance using the sub-haplogroups in addition to macrohaplogroups. We found that sub-haplogroup concordance decreased slightly for MAF>1% (−2.5%) and MAF>0.5% (−0.6%), and only slightly increased using MAF>0.1% (1.4%). Statistical significance is observed between all these comparisons (Table S7a-c). The differences between k_hap_ parameters settings were more pronounced, with all k_hap_ parameter values showing a decrease in concordance (Table S7d-f), likely because all k_hap_ experiments used the Reference Panel filtered at MAF>1%. Larger k_hap_ parameter values performed more poorly than smaller values.

Macrohaplogroup concordance increased only slightly following imputation. There was no statistically significant difference between any of the MAF thresholds, although there was a slight increase in accuracy with decreasing MAF (0.8% to 2.2%, ANOVA p = 0.09). Reference haplotype parameter values from *k*_*hap*_ = 100 to *k*_*hap*_ = 1,000 exhibit minor increases in performance, with larger k_hap_ parameter values leading to relatively poorer imputation performance (Table S8d-f). We note, however, that the mean macrohaplogroup concordance in the ISM data was already >86.7%.

Additionally, we evaluated whether the HaploGrep2 haplogroup quality score improved following imputation. There was no significant difference in haplogroup quality score between MAF thresholds (ANOVA, *p* = 0.56); however, on average there was a small decrease in the quality score (0.6%-0.8%) (Table S9a-c). The parameter values for the number of included reference haplotypes showed statistical differences starting at the contrast *k*_*hap*_ = 100 to *k*_*hap*_ = 1,000, with imputation accuracy decreasing at higher k_hap_ parameter values (Table S9d-f).

Improvements in Haplogroup concordance was also evaluated using Hi-MC to assign haplogroups. Following imputation, there was a mean increase (31.2%-32.5%) in accuracy of haplogroup assignment across different Reference Panel MAF thresholds. However, there was no statistically significant difference between these MAF thresholds (ANOVA, p = 0.83) (Table S10a-c). With an increase in the k_hap_ parameter, a decrease in accurate haplogroup assignment was observed, with contrasts at *k*_*hap*_ = 100 to *k*_*hap*_ = 2,500 becoming statistically significant. These patterns were observed when macrohaplogroups were examined (Table S11a-f). On average, haplogroup concordance ranged from 16.7%-21.0%, while macrohaplogroup concordance ranged from 88.0%-88.4%

Taken together, these findings indicate optimum values of *k*_*hap*_ = 500 for the number of reference haplotypes, and MAF>0.1% for the minor allele frequency threshold of the Reference Panel.

#### Overall Microarray Performance

Using our recommended settings (*k*_*hap*_ = 500, MAF>0.1%), most genotypes were successfully imputed in most cases, with μ_*MCC*_ = 0.618 (95% *confidence interval* [*CI*] = 0.615, 0.620). The best performing chip was the GSA-24v2-0_A1-b37 (*MCC* = 0.658; 95%*CI* = 0.636, 0.681), and the worst performing chip was the HumanOmni2.5S-8v1_B-b37 (*MCC* = 0.381; 95%*CI* = 0.320,0.441) (Table S12).

On average, macrohaplogroups assigned using HaploGrep2.0 from imputed data were concordant with the truth set 88.2% of the time (95%*CI* = 88.1%, 89.4%). The GSAMD-24v2-0_20024620_A1-b37 was the best performing ISM in terms of HaploGrep macrohaplogroup concordance (99.4%; 95%*CI* = 99.2%, 99.7%), while the InfiniumImmunoArray-24v2-0_A-b37 was the worst performing ISM (10.8%; 95%*CI* = 9.6%, 12.0%). On average there was an improvement in concordance of 1.5%. HumanOmni2.5S-8v1_B-b37 had the largest improvement (24.4%). HumanOmni5-4v1_B-b37 was the worst performing ISM, with a 13.6% decrease in concordance (Table S12).

On average, macrohaplogroups assigned using Hi-MC from imputed data were concordant with the truth set 91.8% of the time (95%*CI* = 91.7%, 91.9%). BDCHP-1X10-HUMANHAP240S_11216501_A-b37 was the best performing ISM in terms of Hi-MC macrohaplogroup concordance (99.9%, 95%*CI* = 99.8%, 100%), and InfiniumOmniZhongHua-8v1-3_A1-b37 was the worst performing (28.6%; 95%*CI* = 26.9%, 30.4%). The overall increase in improvement was 24.9% (Table S12), with the HumanOmni5-4v1-1_A-b37 the best performing chip, increasing 43.6%, and HumanOmni1-Quad_v1-0_B-b37 the worst performing, showing a 32.8% decrease in concordance.

#### Overall Haplogroup Concordance

Concordance of individual haplogroups was estimated at the macro-haplogroup level using HaploGrep2.0 and Hi-MC. Before imputation, less than 50% of sequences from macrohaplogroup V were assigned to their connect macrohaplogroup by HaploGrep2.0 (Table S13a), and less than 50% of sequences from macrohaplogroups H, HV, I, M, V, W, X were assigned to their correct macrohaplogroup by Hi-MC (Table S13b). Imputation accuracy as measured by macrohaplogroup concordance using HaploGrep2.0 showed a difference with the ISM dataset ranging from a decrease of 16.6% (HV) to an increase of 52.9% (V). With the exception of L5, all African macrohaplogroups showed a slight decrease (3.12%-0.18%). For the Native American-associated macrohaplogroups, only B showed a decrease (5.02%). Among the East Asian-associated macrohaplogroups, G, N, and Z showed a decrease (0.88%-7.42%). Among the Euro-Indian-associated macrohaplogroups, H, J, and U showed a decrease (0.14%-1.82%). Imputation accuracy as measured by macrohaplogroup concordance using Hi-MC showed a difference with the ISM dataset from a decrease of 15.7% (B) to an increase of 89.9% (M). All African macrohaplogroups showed a slight decrease (8.9%-0.64%). The Native American-associated macrohaplogroups, B and C showed a decrease (0.15%-15.7%). Among the East Asian-associated macrohaplogroups, only N showed a decrease (6.5%). Among the Euro-Indian associated macrohaplogroups, only U showed a decrease (0.8%). However, it should be noted that Hi-MC did not detect any presence of macrohaplogroups F, G, L4, L5, Y, or Z.

### Alzheimer’s Disease Neuroimaging Initiative

We applied MitoImpute to data from 258 participants in the ADNI study, who had provided both WGS [29] and microarray data [17] (Table S14). The ADNI microarray data were mapped to the rCRS and following imputation sites with an IMPUTE2 info score ≤ 0.3 were discarded. Both HaploGrep2 [24] and Hi-MC [25] were used to assign haplogroups to the WGS, microarray, and imputed data. Genotypes were moderately successfully imputed, as measured by MCC (μ_*MCC*_ = 0.322; 95%*CI* = 0.294,0.350). This is in contrast with the ISM for the chip with which ADNI was genotyped (Illumina Human610-Quad BeadChip, Human610-Quadv1_B-b37, μ_*MCC*_ = 0.606; 95%*CI* = 0.576,0.637).

Using HaploGrep2.0, the correct macrohaplogroup to 95.7% of samples for the microarray data, which improved to 97.7% after imputation. Macrohaplogroup V showed any improvement of 66.7%, whereas all other macrohaplogroups showed no change, with the exception of H, which showed a 0.9% decrease (Table S15a). The corresponding improvement using Hi-MC was 37.9% to 95.0%. Macrohaplogroups A, H, J, JT, M, N, V, W, and X all showed improvements, ranging from 27.2% to 100% (Table S15b). The results for macrohaplogroups M, V, W, and X, are particularly noteworthy since they had no correct assignments prior to imputation. Macrohaplogroup HV remained at 0% concordance before and after imputation.

## Discussion

Investigations into the genetic basis of human mitochondrial disease and of evolutionary history rely on the accurate alignment of homologous nucleotide positions, and complete mtDNA sequences [30]. These two factors, in turn, benefit from globally diverse sequences being included in MSAs used in these investigations. The imputation of missing variants can mitigate datasets of incomplete mtSNVs; however, accurate alignment of sequences and consistent placement of gap character states is fraught with difficulty and time consuming for even experienced bioinformaticians [10]. Lack of publicly available reference MSAs and reference panels, therefore, presents a limitation to researchers investigating mitochondrial disease or evolutionary history. We address this limitation by creating a reference MSA from 36,960 globally diverse mtDNA sequences, which was manually curated by experienced researchers to ensure consistency of the placement of gap character states. Aligning novel sequences to our reference alignment will alleviate the pressures of the alignment process by providing a guide for these new sequences.

The reference panel we present here is globally and phylogenetically representative. Despite less than 20% of samples having geographic provenance metadata available, the sample that do contain this information suggest there are at least 103 geographic regions from 49 countries cover all inhabited continents. These include populations usually not represented in major population genetic datasets (e.g., the 1,000 Genomes Project), such as Pacific Islanders and Indigenous Australians. Additionally, all PhyloTree [27] macrohaplogroups are present in our reference alignment and reference panel. To the best of our knowledge, this is the largest and most genetically and geographically diverse curated mtDNA reference panel publicly available. Additionally, as a curated MSA, our reference MSA can be subsampled for use in answering evolutionary and disease-associated research questions. Furthermore, the reference MSA can be used as a reference panel for the imputation of mtSNVs. This reference panel will enable comparison and combined analyses across datasets of differing age and completeness. The reference panel has been packaged into a user-friendly mtSNV imputation pipeline, MitoImpute.

We evaluated how accurately we could impute mtSNVs using our reference panel, as measured by the concordance of assigned haplogroups and Matthews [26] correlation coefficient of genotypes. Across most ISMs, we were able to improve genotype concordance and macrohaplogroup assignment marginally when assigned using HaploGrep2.0 and significantly when using Hi-MC. As HaploGrep2.0 already accurately assigns macrohaplogroups, these results suggest we are successfully imputing phylogenetically informative mtSNVs. Some macrohaplogroups experienced marginal decreases in their correct assignment; however, this does not appear to be biased to any locality outside of Africa. As all haplogroups, except for haplogroups JT and X, experienced an average improvement >30%, this suggests that the reference panel is not biased towards improvement for certain lineages over others. The addition of new sequences to the reference panel will only further increase accurate haplogroup assignment in populations or mtDNA lineages that are still underrepresented. We also tested the practical use of our reference panel by imputing mtSNVs in the ADNI dataset, demonstrating that the reference panel and imputation pipeline can successfully impute genotypes and, in some instances, dramatically increase the correct macrohaplogroup assignment. Given that there are 499 samples in the ADNI genotyping dataset that were not re-sequenced in subsequent phases, this demonstrates the utility of our reference panel for long-term studies that need to bring their older, incomplete dataset to the same standard as newer, complete datasets.

Performance testing of the MitoImpute pipeline using ISM revealed a seemingly counterintuitive result, the decrease in imputation accuracy as the k_hap_ parameter increases. Increasing the k_hap_ parameter increases the number of haplotypes in the reference panel from which IMPUTE2 will impute. We suspect that increasing the number of reference haplotypes beyond 1,000 leads to a greater chance of mismatch between the incomplete sample haplotypes and the reference panel haplotypes, particularly in ISMs with few mtSNVs. Alternatively, highly diverse reference panels may contain a large number of haplotypes uninformative for imputing variants missing from the study sample, which has previously been noted by [31]. The limitations of the MAF and k_hap_ parameters, we suspect, is due to a dearth of mtSNVs in some ISMs. Datasets with a small number of variants from which to impute missing mtSNVs will always present this limitation, and we recommend users proceed with caution when using these datasets for subsequent analyses.

Our reference panel provides an opportunity for datasets with limited mitochondrial genetic variation to be analysed with a more complete set of genetic variants and a more accurate assignment of haplogroups. The global disparity in medical research is evident in the high proportion of European individuals (~78%) association study catalogues [32]. The 1,000 Genomes Project phase 3 includes 2,504 individuals from 26 populations, however, these individuals were often sampled from 1-3 cities within geographically diverse countries, such as China. Our reference panel contains sequences from at least 103 regions in at least 49 countries, capturing a more globally-representative sample of mitochondrial genetic diversity. The diversity included in our reference panel will allow researchers to perform imputation in under-represented human populations, contributing to solving the disparity in medical genomics research. This study also highlights the imperative to include accurate and detailed metadata when submitting sequences to public repositories, such as GenBank. Having only geographic provenance metadata available for 19.3% of downloaded GenBank sequences limits our ability to determine regions underrepresented in DNA databases. As haplogroups are only useful for determining geographic provenance at a fine sub-haplogroup level [1], haplogroups cannot be relied on as geographic proxies.

## Availability of source code

Project name: MitoImpute

Project home page: https://github.com/sjfandrews/MitoImpute

Operating system(s): Linux; OSX

Programming language: Python

Other requirements: Snakemake 5.30.1

License: MIT License

## Availability of supporting data

The data set(s) supporting the results of this article is(are) available in the in MitoImpute GitHub repository, (10.5281/zenodo.4338785).

## Abbreviations

ADNI: Alzheimer’s disease neuroimaging initiative
ISMs: in silico microarrays
MAF: Minor allele frequency
MCC: Matthews Correlation Coefficient
MSA: multiple sequence alignment
mtDNA: mitochondrial DNA
mtSNVs: mitochondrial DNA single nucleotide variants
WGS: whole-genome sequence

## Competing interests

AMG served on the scientific advisory board for Denali Therapeutics from 2015-2018. She has also served as a consultant for Biogen, AbbVie, Pfizer, GSK, Eisai and Illumina

## Funding

Dr. Judy Pa and Christopher Patterson were supported by the National Institute on Aging (R01AG054617 PI: Judy Pa). BFH and AMG are supported by the JPB Foundation (http://www.jpbfoundation.org). RHS is supported by P30 AG035982. SJA is supported by the JPB Foundation and the Alzheimer’s Association (AARF-20-675804).

## Author’s Contributions

TWM, SJA conceptualized, designed the study and coordinated data collection, development of software code, and carried out initial analyses and drafted the initial manuscript. BFH, CP and DP were involved in implementation of the computer code and supporting algorithms and reviewed the manuscript. HP, JP, RHS, AG, SE and LSJ provided mentorship and reviewed the manuscript. All authors approved the final manuscript as submitted and agree to be accountable for all aspects of the work.

## Acknowledgments

Data collection and sharing for this project was funded by the Alzheimer’s Disease Neuroimaging Initiative (ADNI) (National Institutes of Health Grant U01 AG024904) and DOD ADNI (Department of Defense award number W81XWH-12-2-0012). ADNI is funded by the National Institute on Aging, the National Institute of Biomedical Imaging and Bioengineering, and through generous contributions from the following: AbbVie, Alzheimer’s Association; Alzheimer’s Drug Discovery Foundation; Araclon Biotech; BioClinica, Inc.; Biogen; Bristol-Myers Squibb Company; CereSpir, Inc.; Cogstate; Eisai Inc.; Elan Pharmaceuticals, Inc.; Eli Lilly and Company; EuroImmun; F. Hoffmann-La Roche Ltd and its affiliated company Genentech, Inc.; Fujirebio; GE Healthcare; IXICO Ltd.; Janssen Alzheimer Immunotherapy Research & Development, LLC.; Johnson & Johnson Pharmaceutical Research & Development LLC.; Lumosity; Lundbeck; Merck & Co., Inc.; Meso Scale Diagnostics, LLC.; NeuroRx Research; Neurotrack Technologies; Novartis Pharmaceuticals Corporation; Pfizer Inc.; Piramal Imaging; Servier; Takeda Pharmaceutical Company; and Transition Therapeutics. The Canadian Institutes of Health Research is providing funds to support ADNI clinical sites in Canada. Private sector contributions are facilitated by the Foundation for the National Institutes of Health (www.fnih.org). The grantee organization is the Northern California Institute for Research and Education, and the study is coordinated by the Alzheimer’s Therapeutic Research Institute at the University of Southern California. ADNI data are disseminated by the Laboratory for Neuro Imaging at the University of Southern California.

## Supplementary Information

### Supplementary Methods

The following search term was used to identify whole human mtDNA sequences from GenBank on 2018-07-18:

(016500[SLEN]:016600[SLEN]) AND Homo[Organism] AND mitochondrion[FILT] AND complete genome NOT (Homo sp. Altai OR Denisova hominin OR neanderthalensis OR heidelbergensis OR consensus OR ancient human remains OR shotgun)

Reference Alignment

We used publicly available PhyloTree (van Oven and Kayser, 2009) sequences to create a large (n=7,747) reference alignment with the revised Cambridge Reference Sequence (rCRS) (Andrews et al., 1999) site numbering convention. Inclusion of rCRS in the reference alignment ensures that site numbering conventions are maintained and verified as new sequences are added. We aligned sequences in batches of 50 using the L-INS-i version of MAFFT (Katoh and Standley, 2013), then combined the batches, resolving inconsistent gap placements manually. rCRS site numbers were preserved by removing sites at which gaps were introduced in the rCRS during the alignment process.

### Supplementary Tables

Table S1: Sequences included in the Reference Panel alignment

Table S2: Strand files downloaded from the Wellcome Centre

Table S3: Variable sites found in the Reference Panels at varying minor allele frequencies (MAF) 1%, 0.5%, and 0.1%

Table S4: Summary table of geographic provenance of samples in the reference alignment and panel extracted from GenBank metadata.

Table S5a-f: MCC genotype imputation accuracy across MAF and k_hap_ settings

Table S6a-f: IMPUTE2 info score across MAF and k_hap_ settings

Table S7a-f: HaploGrep2.0 haplogroup concordance across MAF and k_hap_ settings

Table S8a-f: HaploGrep2.0 macrohaplogroup concordance across MAF and k_hap_ settings

Table S9a-f: HaploGrep2.0 haplogroup quality score across MAF and k_hap_ settings

Table S10a-f: Hi-MC haplogroup concordance across MAF and k_hap_ settings

Table S11a-f: Hi-MC macrohaplogroup concordance across MAF and k_hap_ settings

Table S12: Per-chip performance summary using recommended parameter settings (MAF>0.1% and k_hap=500)

Table S13a-b: Proportion of macrohaplogroups correctly assigned using HaploGrep2.0 and Hi-MC before and after imputation

Table S14: ADNI samples with genotype and whole genome sequencing data

Table S15a-b: Macro-haplogroup concordance between genotyped and imputed ADNI data using HaploGrep2.0 and Hi-MC

## Notes

https://github.com/sjfandrews/MitoImpute

### Summary of Updates

Updated results and interpretation

## References

1. Underhill PA and Kivisild T. Use of Y Chromosome and Mitochondrial DNA Population Structure in Tracing Human Migrations. Annual Review of Genetics. 2007;41 1:539–64. doi:https://doi.org/10.1146/annurev.genet.41.110306.130407.

2. Gorman GS, Chinnery PF, DiMauro S, Hirano M, Koga Y, McFarland R, et al. Mitochondrial diseases. Nature Reviews Disease Primers. 2016;2:16080. doi:https://doi.org/10.1038/nrdp.2016.80.

3. Gonçalves VF, Giamberardino SN, Crowley JJ, Vawter MP, Saxena R, Bulik CM, et al. Examining the role of common and rare mitochondrial variants in schizophrenia. PLoS One. 2018;13 1:e0191153. doi:https://doi.org/10.1371/journal.pone.0191153.

4. Huang J, Howie B, McCarthy S, Memari Y, Walter K, Min JL, et al. Improved imputation of low-frequency and rare variants using the UK10K haplotype reference panel. Nature Communications. 2015;6:8111. doi:https://doi.org/10.1038/ncomms9111.

5. Yoo S-K, Kim C-U, Kim HL, Kim S, Shin J-Y, Kim N, et al. NARD: whole-genome reference panel of 1779 Northeast Asians improves imputation accuracy of rare and low-frequency variants. Genome Medicine. 2019;11 1:64. doi:https://doi.org/10.1186/s13073-019-0677-z.

6. Sariya S, Lee JH, Mayeux R, Vardarajan BN, Reyes-Dumeyer D, Manly JJ, et al. Rare Variants Imputation in Admixed Populations: Comparison Across Reference Panels and Bioinformatics Tools. Frontiers in Genetics. 2019;10:239. doi:https://doi.org/10.3389/fgene.2019.00239.

7. Das S, Forer L, Schonherr S, Sidore C, Locke AE, Kwong A, et al. Next-generation genotype imputation service and methods. Nature Genetics. 2016;48 10:1284–7. doi:https://doi.org/10.1038/ng.3656.

8. Zheng H-F, Ladouceur M, Greenwood CMT and Richards JB. Effect of genome-wide genotyping and reference panels on rare variants imputation. Journal of Genetics and Genomics. 2012;39 10:545–50. doi:https://doi.org/10.1016/j.jgg.2012.07.002.

9. Browning BL and Browning SR. A unified approach to genotype imputation and haplotype-phase inference for large data sets of trios and unrelated individuals. American Journal of Human Genetics. 2009;84 2:210–23. doi:https://doi.org/10.1016/j.ajhg.2009.01.005.

10. Golubchik T, Wise MJ, Easteal S and Jermiin LS. Mind the gaps: evidence of bias in estimates of multiple sequence alignments. Molecular Biology and Evolution. 2007;24 11:2433–42. doi:https://doi.org/10.1093/molbev/msm176.

11. Morrison DA. Why Would Phylogeneticists Ignore Computerized Sequence Alignment? Systematic Biology. 2009;58 1:150–8. doi:https://doi.org/10.1093/sysbio/syp009.

12. Morrison DA. Is Sequence Alignment an Art or a Science? Systematic Botany. 2015;40 1:14–26. doi:https://doi.org/10.1600/036364415X686305.

13. The 1000 Genomes Project Consortium. A global reference for human genetic variation. Nature. 2015;526 7571:68–74. doi:https://doi.org/10.1038/nature15393.

14. McCarthy S, Das S, Kretzschmar W, Delaneau O, Wood AR, Teumer A, et al. A reference panel of 64,976 haplotypes for genotype imputation. Nature Genetics. 2016;48 10:1279–83. doi:https://doi.org/10.1038/ng.3643.

15. Hudson G, Gomez-Duran A, Wilson IJ and Chinnery PF. Recent Mitochondrial DNA Mutations Increase the Risk of Developing Common Late-Onset Human Diseases. PLOS Genetics. 2014;10 5:e1004369. doi:https://doi.org/10.1371/journal.pgen.1004369.

16. Howie BN, Donnelly P and Marchini J. A flexible and accurate genotype imputation method for the next generation of genome-wide association studies. PLoS Genetics. 2009;5 6:e1000529. doi:https://doi.org/10.1371/journal.pgen.1000529.

17. Saykin AJ, Shen L, Foroud TM, Potkin SG, Swaminathan S, Kim S, et al. Alzheimer’s Disease Neuroimaging Initiative biomarkers as quantitative phenotypes: Genetics core aims, progress, and plans. Alzheimers Dement. 2010;6 3:265–73. doi:https://doi.org/10.1016/j.jalz.2010.03.013.

18. Lott MT, Leipzig JN, Derbeneva O, Xie HM, Chalkia D, Sarmady M, et al. mtDNA Variation and Analysis Using Mitomap and Mitomaster. Current Protocols in Bioinformatics. 2013;44:1 23 1–6. doi:https://doi.org/10.1002/0471250953.bi0123s44.

19. Katoh K and Standley DM. MAFFT Multiple Sequence Alignment Software Version 7: Improvements in Performance and Usability. Molecular Biology and Evolution. 2013;30 4:772–80. doi:https://doi.org/10.1093/molbev/mst010s.

20. Kearse M, Moir R, Wilson A, Stones-Havas S, Cheung M, Sturrock S, et al. Geneious Basic: An integrated and extendable desktop software platform for the organization and analysis of sequence data. Bioinformatics. 2012;28 12:1647–9. doi:https://doi.org/10.1093/bioinformatics/bts199.

21. Andrews RM, Kubacka I, Chinnery PF, Lightowlers RN, Turnbull DM and Howell N. Reanalysis and revision of the Cambridge reference sequence for human mitochondrial DNA. Nature Genetics. 1999;23 2:147-. doi:https://doi.org/10.1038/13779.

22. Wong TKF, Kalyaanamoorthy S, Meusemann K, Yeates DK, Misof B and Jermiin LS. A minimum reporting standard for multiple sequence alignments. NAR Genomics and Bioinformatics. 2020;2 2 doi:https://doi.org/10.1093/nargab/lqaa024.

23. Rayner W: Genotyping chips strand and build files. https://www.well.ox.ac.uk/~wrayner/strand/. Accessed 28 May 2018.

24. Weissensteiner H, Pacher D, Kloss-Brandstätter A, Forer L, Specht G, Bandelt H-J, et al. HaploGrep 2: mitochondrial haplogroup classification in the era of high-throughput sequencing. Nucleic Acids Research. 2016;44 W1:W58–W63. doi:https://doi.org/10.1093/nar/gkw233.

25. Smieszek S, Mitchell SL, Farber-Eger EH, Veatch OJ, Wheeler NR, Goodloe RJ, et al. Hi-MC: a novel method for high-throughput mitochondrial haplogroup classification. PeerJ. 2018;6:e5149. doi:https://doi.org/10.7717/peerj.5149.

26. Matthews BW. Comparison of the predicted and observed secondary structure of T4 phage lysozyme. Biochimica et Biophysica Acta (BBA) - Protein Structure. 1975;405 2:442–51. doi:https://doi.org/10.1016/0005-2795(75)90109-9.

27. van Oven M. PhyloTree Build 17: Growing the human mitochondrial DNA tree. Forensic Science International: Genetics Supplement Series. 2015;5:e392–e4. doi:https://doi.org/10.1016/j.fsigss.2015.09.155.

28. Köster J and Rahmann S. Snakemake – a scalable bioinformatics workflow engine. Bioinformatics. 2012;28 19:2520–2. doi:https://doi.org/10.1093/bioinformatics/bts480.

29. Ridge PG, Wadsworth ME, Miller JB, Saykin AJ, Green RC and Kauwe JSK. Assembly of 809 whole mitochondrial genomes with clinical, imaging, and fluid biomarker phenotyping. Alzheimer’s & Dementia. 2018;14 4:514–9. doi:https://doi.org/10.1016/j.jalz.2017.11.013.

30. Kumar S and Filipski A. Multiple sequence alignment: in pursuit of homologous DNA positions. Genome Research. 2007;17 2:127–35. doi:https://doi.org/10.1101/gr.5232407.

31. Nelson SC, Stilp AM, Papanicolaou GJ, Taylor KD, Rotter JI, Thornton TA, et al. Improved imputation accuracy in Hispanic/Latino populations with larger and more diverse reference panels: applications in the Hispanic Community Health Study/Study of Latinos (HCHS/SOL). Human Molecular Genetics. 2016;25 15:3245–54. doi:https://doi.org/10.1093/hmg/ddw174.

32. Sirugo G, Williams SM and Tishkoff SA. The Missing Diversity in Human Genetic Studies. Cell. 2019;177 1:26–31. doi:https://doi.org/10.1016/j.cell.2019.02.048.

